# Dynamic Dark Root Chamber – Advancing non-invasive phenotyping of roots kept in darkness using infrared imaging

**DOI:** 10.1101/2024.02.16.580252

**Authors:** Simon Pree, Ivan Kashkan, Katarzyna Retzer

**Affiliations:** Department of Forest and Soil Sciences, Institute of Forest Ecology, University of Natural Resources and Life Sciences (BOKU), Peter-Jordan Straße 82, Vienna, 1190, Austria; Laboratory of Hormonal Regulations in Plants, Institute of Experimental Botany, Czech Academy of Sciences, Prague, Czechia; PSI (Photon Systems Instruments), spol. s.r.o., 66424 Drásov, Czech Republic; Imaging Facility of Institute of Experimental Botany AS CR, Prague, Czech Republic

**Keywords:** dynamic root growth phenotyping, seminal roots, root growth traits, root imaging, Raspberry Pi, infrared LED, dark grown roots, root system architecture

## Abstract

Root phenotyping is a challenging task that would require monitoring root growth in soil under dark conditions to mimic natural conditions, while allowing the shoot to grow in light. Most existing methods involve exposing the roots to light, which substantially alters their growth and function. In this paper, we present an improved imaging system that can overcome this limitation of experiments performed in laboratories. The Dynamic Dark Root imaging Chamber (DDrC) enables continuous monitoring and image acquisition to track the dynamic development of root architecture under controlled growth conditions. Our imaging system is based on a Raspberry Pi camera module and infrared LEDs, which do not induce any stress responses in the roots. The DDrC setup is simple, affordable, and suitable for dynamic phenotyping experiments. We provide a detailed tutorial for the assembly and adjustment of the imaging chamber. We conclude that our system is a valuable tool for studying the genetic and environmental factors that affect the root system architecture and development, and for identifying the root traits that are related to plant adaptation and performance.

## 1. Introduction

The root is a crucial organ that anchors the plant in the soil and facilitates water and nutrient uptake which is essential for plant growth and survival (Pierik and Testerink, 2014; Retzer and Weckwerth, 2021, 2023; Pree *et al*., 2023). While it is relatively easy to measure and analyze the shoot phenotype of plants in the field or in the laboratory under physiological relevant conditions, it is still very difficult, time-consuming, or harmful for the root to obtain dynamic data of the whole root system (York, Griffiths and Maaz, 2022).

Roots have evolved to grow vertically downward in the soil by sensing and responding to gravity and light signals (Wyatt and Kiss, 2013; Retzer and Weckwerth, 2023). However, recent studies have revealed that exposing roots to light not only influences their direction of growth, but also interferes with other root responses, altering their growth rate and function compared to roots grown in the dark (Xu *et al*., 2013; Silva-Navas *et al*., 2015; Shi *et al*., 2018; García-González, Lacek and Retzer, 2021).

Root phenotyping experiments involve three steps: (a) experimental design, which includes plant growth and sometimes stress application, (b) image acquisition, and (c) image segmentation and data analysis. While many publications suggest advanced methods of data analysis using deep learning techniques, the experimental designs and image acquisition frequently expose the root to light, which can affect the root phenotype. The development of reliable, automated image processing software is crucial to evaluate a large number of individual roots, due to their high variability, and to associate different phenotypes with specific quantitative trait loci (QTLs) (Fernandez *et al*., 2022; LaRue *et al*., 2022; Lube *et al*., 2022; Ohlsson *et al*., 2023). The extraction and analysis of multiple root traits and growth dynamics are constantly improved (Seethepalli *et al*., 2021).

Our current research involves to monitor various characteristics of barley’s seminal roots in soil. Seminal roots are the initial roots that emerge during germination, they typically range from 4 to 8 in number and explore soil water to grow towards water and nutrient abundant areas (Richards, Watt and Rebetzke, 2007). They allow the plant to establish quickly and securely in soil. The number of seminal roots, as well as their angle and speed of growth through the soil, are vital for cereal performance (Richards, Watt and Rebetzke, 2007). The root opening angle and the growth rate influence the depth and spreading of the root system. Several genes have been linked to the regulation of root opening angle and a narrow root system is correlated with deep-rooted plants in dry regions, while wide angles are found in varieties adapted to areas with sufficient rainfall (Morris *et al*., 2017; Salvi, 2017; Maqbool *et al*., 2022). The number of seminal roots is proposed to vary according to the nutrient and overall resource availability. The root growth rate is flexible and strongly affected by the soil properties. The elongation rates of seminal roots are crucial for our studies as they indicate the roots’ ability to quickly spread in the soil and access deeper soil layers (Maqbool *et al*., 2022). Therefore, the proposed method offers a significant advantage in tracking the seminal expansion rate continuously and allows in a simple way under more natural growth conditions to examine a high number of different genotypes and growth conditions in standard growth chambers.

Summarized, we describe a simple and affordable imaging system for young root systems, which can be used to monitor root outgrowth from germination, on agar plates, germination paper or soil. The roots are grown in darkness, and the shoot emerges from the imaging chamber to be illuminated by the light sources available in the growth room or chamber. Root images are captured with a Raspberry Pi camera at any intervals needed to study the desired root traits. For illumination during image taking infrared LEDs with a wavelength of 880nm are used, which were repetitively reported to not induce measurable stress responses of roots.

## 2. The negative impact of direct illumination on root growth and function requires a dark grown root setup

Plants are highly responsive to their environment and adjust their growth and development accordingly. One of the most important environmental factors that plants perceive and respond to is light. Different wavelengths of light spectrum can provide energy for photosynthesis, or act as a signal that informs plants about the quality of the light source (Miotto *et al*., 2021; van Gelderen *et al*., 2021). Plants express various photoreceptors that sense different wavelengths of light and accordingly trigger specific responses in the shoot, but also in the root (Van Gelderen, Kang and Pierik, 2018; Retzer and Weckwerth, 2021, 2023; Lopez *et al*., 2023). While shoots are directly exposed to light, roots are usually shielded from direct illumination by the soil. However, roots can still sense light indirectly through signals from the shoot or through light penetration in the soil. Root illumination can have detrimental effects on root growth, architecture and function, as well as on the overall plant performance and fitness. How root illumination affects plant fitness is an emerging research area that has gained more attention in recent years (Xu *et al*., 2013; Silva-Navas *et al*., 2015; García-González, Lacek and Retzer, 2021; Lacek *et al*., 2021; Cabrera, Conesa and Del Pozo, 2022; González-García *et al*., 2022).

Currently, one of the most extensively studied model plants for root illumination is *Arabidopsis thaliana*, but also crop species, especially barley, are recently studied more extensively in the so-called D-root setup. Silva-Navas et al., 2015 and 2016, showed in detailed studies the impact of direct illumination on *Arabidopsis thaliana* root growth, function and stress responses, which also results in delimited shoot growth (Silva-Navas *et al*., 2015, 2016). Root meristem activity, which determines the root growth potential, decreases when roots are exposed to light, while the elongation rate, which determines the root length, increases and directional growth deviates more from vertical upon direct root illumination (Mo *et al*., 2015; Silva-Navas *et al*., 2016; Shi *et al*., 2018; Lacek *et al*., 2021). These changes in root growth are accompanied by changes in cell morphology, such as altered cell size, as well as altered cytoskeleton arrangement, which are well described processes that steer root growth modulation (Dyachok *et al*., 2011; Silva-Navas *et al*., 2016; Halat, Gyte and Wasteneys, 2020; García-González and van Gelderen, 2021). Light stress also affects the hormonal balance and responses in the root. Among others, direct root illumination alters the responses to exogenously applied hormones, such as auxin, cytokinin, ethylene, and abscisic acid, as well as influences the endogenous hormone homeostasis, especially of auxin and cytokinin (Silva-Navas *et al*., 2015; Miotto *et al*., 2021; Cabrera, Conesa and Del Pozo, 2022). Those phytohormones are important regulators of root development, especially of lateral root (LR) formation, which determines the root system architecture. Root illumination reduces LR outgrowth, where blue and UV receptors, are crucial for the inhibition (Silva-Navas *et al*., 2015; Yang and Liu, 2020).

Root illumination also affects the nutrient uptake and translocation in the plant, which are essential for plant growth and survival. Illumination of the roots alters uptake efficiency and responses to nutrient deficiency in the root, as well as the accumulation of nutrients in the shoot (Silva-Navas *et al*., 2015, 2019). Root illumination also enhances the responses to additional stress, such as salt, drought, and heat stress, by altering the expression of stress-responsive genes and the accumulation of stress-related metabolites (Silva-Navas *et al*., 2015, 2019; Miotto *et al*., 2021).

Root illumination can also have an impact on the metabolome of the plant, especially on the production of secondary metabolites that have medicinal or nutritional value. Direct root illumination increased the accumulation of artemisinin in the shoot of *Artemisia annua* and hypericin in the shoot of *Hypericum perforatum*, two important phytochemicals. The authors suggest that root illumination induces a stress response in the plant that triggers the biosynthesis of these compounds (Paponov, Ziegler and Paponov, 2023). Another interesting effect of root illumination is the increased root hair outgrowth rate and length near the elongation zone of *Arabidopsis thaliana* roots, which is believed to occur due to disbalanced ratios between reactive oxygen species and their scavengers, but further studies are required to understand this response (Yokawa, Koshiba and Baluška, 2014; Silva-Navas *et al*., 2016; Yokawa, Kagenishi and Baluška, 2016; García-González, Lacek and Retzer, 2021). García-González et al., 2021 further showed that the enhancement of root hair outgrowth is higher when sucrose is added to the growth medium, suggesting a synergistic effect of light and sugar on root hair development (García-González, Lacek and Retzer, 2021).

Besides Arabidopsis, other plant species also show responses to direct root illumination. Shi et al. 2018 showed that rapeseed, maize and barley exhibit strong light avoidance growth under short-term illumination. They further showed that IR LEDs do not affect root growth and orientation, suggesting that IR light does not induce stress responses in the root (Shi *et al*., 2018). In their long-term experiments, root volume was reduced in the root side closer the light source, which corresponds to the previously described influence on root architecture by direct illumination (Xu *et al*., 2013; Silva-Navas *et al*., 2015; Shi *et al*., 2018).

Despite the known negative impact of growing roots exposed directly to light on plant fitness, most lab experiments with plants are performed with roots fully exposed to light. This can lead to artefacts and outcomes that do not reflect the true responses of plants under natural growth conditions. Considering all the aforementioned factors, there arises a need for an imaging system capable of accurately describing root architecture without exposing the roots to disturbing light irradiation, while ensuring shoot growth in light. In order to be widely accepted, such a system should be as affordable and straightforward as traditional laboratory plant growing methods. For colleagues who want to track root growth continuously in a controlled environment, we suggest the simple and affordable IR LED-based approach. Our approach uses IR LEDs (880nm) to shortly illuminate the roots for image acquisition, while avoiding the stress effects of other light wavelengths (Silva-Navas *et al*., 2015; Shi *et al*., 2018). IR LEDs seem to have no obvious effect on root fitness (Wells *et al*., 2012; Shi *et al*., 2018; Dermendjiev *et al*., 2023). We were previously using IR LEDs to study growth dynamics of etiolated seedlings and we couldn’t distinguish growth behaviour from control plants in fully etiolated growth conditions (Kashkan, Garcia-González, *et al*., 2022; Kashkan, Hrtyan, *et al*., 2022). This approach can be used to monitor root outgrowth from germination, on agar plates, germination paper or soil, and to study various dynamic root traits and responses.

## 3. Assembly of the Dynamical Dark Root imaging Chamber (DDrC)

The imaging chamber consists of a corpus and various electronic components. It depends on the preferences and scientific aims of the individual scientist (as described below), which material is used for the corpus, and how big the chamber shall be. Also, it depends on the investigated plant species and type of treatment if seedlings are germinated and monitored grown in agar plates, soil-filled rhizoboxes or on germination paper. Our research interest is currently focused on barley seminal root outgrowth, therefore we focus during this manual on how to assemble a chamber with dimensions required to image barley seedling germination and seminal root outgrowth in a soil-filled self-made rhizobox, which was already described in Lube et al., 2022 (Lube *et al*., 2022). Therefore, we are using modified polystyrene plates, cut at the top to allow the shoot to grow out of the imaging chamber. This setup enables us to keep the dimensions small enough to place several imaging chambers next to each other on shelves overall used in growth chambers with the average height known to track *Arabidopsis thaliana* seedlings grown in standard agar plates. We provide a detailed list of items, including their costs, required to build up your own imaging chamber and describe all steps needed to assemble the chamber.

### 3.1. Selection of the material for the imaging chamber corpus

In this manuscript we present a method for constructing an imaging chamber that enables the observation of the growth characteristics of barley seminal roots, including their growth rate, root opening angle, and number. In this setup the matching rhizobox can accommodate up to five barley seedlings and monitor their root development from the start of germination till one week.

For image acquisition and camera control, components from the Raspberry Pi ecosystem were utilized due to their accessibility and low costs. At the time of publishing of this preprint the parts could be sourced for less than200€.

The size of the chamber can be adjusted according to personal research interests. The chamber is adaptable to different plant species and experimental durations, depending on the research objectives. The key factors for re-designing the chamber are the placement of the camera and the distance of the specimen from the camera, which determine the quality and accuracy of the root images. We will later further describe the proper position of the LEDs, which needs to be altered depending on the implementation of agar plates or rhizoboxes filled with soil.

The material of the chamber can be altered according to personal interests, but it should be opaque and not accumulate heat, to avoid interference with root growth and image acquisition. If the material tends to heat up, we recommend adding fans to cool down the chamber. In this manuscript, we use solid cardboard for the corpus of the chamber, because it is inexpensive, easy to manipulate, and proven to be effective in previous imaging setups (Kashkan, Garcia-González, *et al*., 2022; Kashkan, Hrtyan, *et al*., 2022).

The chamber that we describe in this manuscript has the dimensions of 30 x 24 x

9.5 cm and is suitable for tracking the root growth of *Arabidopsis thaliana* seedlings from germination to about two weeks, or barley seedlings for one week (Fig. 1). The growth rate of the seedlings may vary according to the temperature in the growth chamber and illumination setups of the shoot. The size of the described chamber allows placement of multiple quantities next to each other on standard shelves of regular growth rooms.

**Figure 1:**
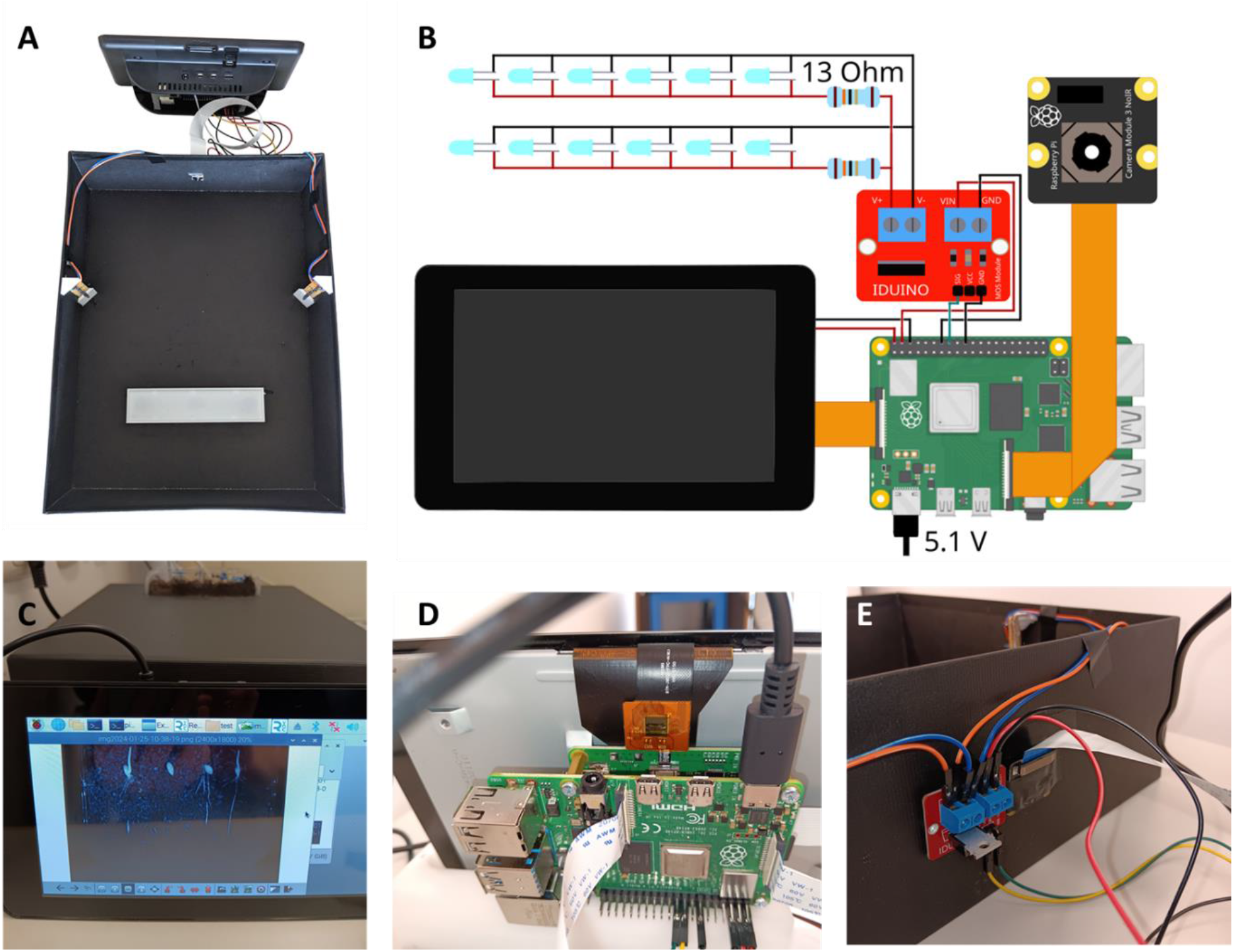
The displayed imaging chamber, powered by the Raspberry Pi and equipped with IR LEDs, enables precise monitoring and analysis of dynamic plant growth and root development. A) The picture provides an overview of the essential components within the imaging chamber. The corpus as structural framework houses the entire setup, and can be individually adjusted regarding its material and size. The infrared (IR) LEDs emit light at 880nm, which allow to acquire images of roots shaded from white light without influencing their physiology. The integrated camera is positioned within the corpus opposite the specimen. The Raspberry Pi is placed outside the imaging chamber and serves as the computational hub for data processing and control. B) Detailed component scheme of the individual components and their interconnections. It illustrates how the camera, IR LEDs, and Raspberry Pi are electrically connected and indicates the physical arrangement of these components within the chamber. The detailed list of the components and their features is available in the main text of the manuscript. C) In this view, the imaging chamber is closed, enclosing the soil-filled petri-dish used as mini-rhizobox. The rhizobox is placed such that the emerging coleoptile will grow out of the chamber, which allows its illumination in any growth chamber as usual. At the front of the picture the display of the Raspberry Pi is visible, which shows an image acquired from the roots. D) The picture displays the internal components of the Raspberry Pi computer module. E) The picture provides a close-up on the camera module that is inserted into the corpus. It further shows the cables necessary for connecting the camera module and IR LEDs to the Raspberry Pi, including the meticulous wiring required for seamless operation.

### 3.2. Assembly of the individual parts

Material required to build up one chamber and costs per item:

- Raspberry Pi 4, 1 GB RAM – 40€
- Raspberry Pi Camera 3 NoIR, 76^°^ - 30€
- Raspberry Pi 7^”^ Touch Screen Display – 70€
- case for Display and Raspberry Pi – 13€
- SanDisk Ultra microSDHC 32 GB - 6€
- Raspberry Pi USB-C power supply 5,1V / 3,0A – 8€
- Iduino ST1168 transistor module 2.50€
- 12^*^Kingbright Infrared LED 880 nm L-7104SF4BT – 6€
- 2 ^*^13 Ohm resistor – 0.20€ strip board 10 cm – 1€
- 6 Dupont connector cables ∼0.20€
- corpus – in this manuscript black cardboard box 30x24x9.5cm - 5€
- optional: USB Stick – 10€
- optional: 3D printed mounts for the LEDs, support for the plates: 0.50€

For image acquisition we use the Raspberry Pi Camera 3 NoIR (no infrared filter), which is mounted in the centre of the wall of the box, at a distance of 21 cm, facing the specimen (Fig. 1 A and E). In case the dimensions of the chambers need to be altered, the camera should be positioned in a way that its field of view would cover the entire plate or rhizobox where the roots are growing (Fig. 1 C).

The camera is connected to a Raspberry Pi 4 single board computer with 1 GB RAM, which is sufficient to control the camera module (Fig. 1 D). If image post-processing is planned to be done directly on the Raspberry Pi 4 one might want to get a model with more RAM or the Raspberry Pi 5 that possess a more capable processor. The images can be stored on the microSD card in the Raspberry Pi 4 or a USB stick. Alternatively, if infrastructure permits, a server for remote access and data management can be used.

To obtain pictures we mounted o6 infrared LEDs on each side wall of the box, which are facing the direction of the plate/rhizobox at a 45° angle (Fig. 1 A). In order to avoid reflections on the surface of the soil-filled rhizobox, which would disturb the image analysis later on, the position of the LEDs is important. For the setup described here, the LEDs are mounted at a distance of 10cm to the plate. The LEDs are switched in parallel via a 13 Ohm resistor. As it is not recommended to use the GPIO pins for high output, the LEDs are connected to a MOSFET transistor module, which is controlled by the GPIO pins and powered with the 5V pins of the Raspberry Pi 4 (Fig. 1 B). To ensure smooth operation of the imaging chamber we attached a 7-inch touchscreen display to the Raspberry Pi 4 (Fig. 1 C). The screen is used to start and stop the experiment and set basic parameters including experiment name, time between image acquisition and distance from the camera to the object. To track the progress of the experiment over time we chose to display the elapsed time while the experiment is running.

### 3.3. Operating system and code

Raspberry Pi OS was installed on the microSD card as the operating system. Due to issues with the touchscreen on the standard version we choose Raspberry Pi OS Legacy 64 bit with desktop was, which is based on Debian 12 and uses the Linux kernel version 6.1 (Fig. 1). It can be downloaded at https://www.raspberrypi.com/software/operating-systems/.

We wrote the script for the graphical user interface and image acquisition in Python 3. It includes the code for a graphical user interface that allows to control the experiment and a loop that transfers and saves images at the predefined time. For image preview and image acquisition we applied the modules called ‘libcamera’ and ‘picamera 2’. While it is planned to make the script available on Github in the future, for now we can provide it via email on request.

### 3.4. Imaging settings

The frequency of image acquisition can be selected with a slider on the experiment settings window. The IR LEDs are only switched on for 2 seconds during image acquisition. Exposure is set to a fixed time and saturation was set to 0 for greyscale images. Images are taken at the maximum resolution of the Raspberry Pi camera module 3, which is 4608 x 2592. For our experiment setup using plates we crop the images to a size of 2400 x 1800 pixel in order to exclude the sections that don’t contain the research object. As a file format .png was selected and the image file name contains information of the date and time.

## 4. Post-processing of obtained images

In our setup with the Raspberry Pi v3 camera module, which obtains images in a 72° angle, no distortion effects are observable, and we don’t include post-processing steps in our protocol. But we advise, if another camera module with a wider angle is used to add a step to include correction of the fisheye effect. Additionally, to enhance image contrast for potentially improved segmentation results during image analysis, it may be beneficial to normalize the obtained greyscale images. This approach is recommended if the variations in shades of a colour – in this case, grey – are more important than their absolute values (Depeursinge, Fageot and Al-Kadi, 2017). The images in Fig. 2 A show the dynamics of barley root growth dynamics from the moment of germination and additionally the emergence of the coleoptile. Four images demonstrating the germination process in the first 48 h were selected. The imaging chamber facilitates the evaluation of several traits, such as root opening angle, seminal root number, root growth rate, mesocotyl growth rate, and adaptation to different soil textures and compositions. Root growth rate acquisition and evaluation of the primary root compared to the first pair of seminal roots, as well the root opening angle establishment over time with ImageJ/Fiji is described and visualized in the Suppl. Fig. 1. The images are further suitable for segmentation with the RhizoVision Explorer (Seethepalli et al., 2021), a software tool that automatically extracts root traits from digital images (Fig. 2 B). Maximum root width and convex hull area expansion over time is easily to obtain from the obtained images (Suppl. Fig. 2). It should be considered that the segmentation process is more efficient when the soil is dark and humidity levels in the rhizobox are low, which results in high contrast levels between the barley root and the background. The imaging setup allows continuous monitoring of the root growth status of the germinating seedlings, also over longer time periods, as the barley shoot is able to grow out of the imaging chamber (Fig. 2 C and D). The touch screen enables the selection of already taken images, which allows qualitative control of the experimental setup during the running experiment (Fig. 2 E).

**Figure 2:**
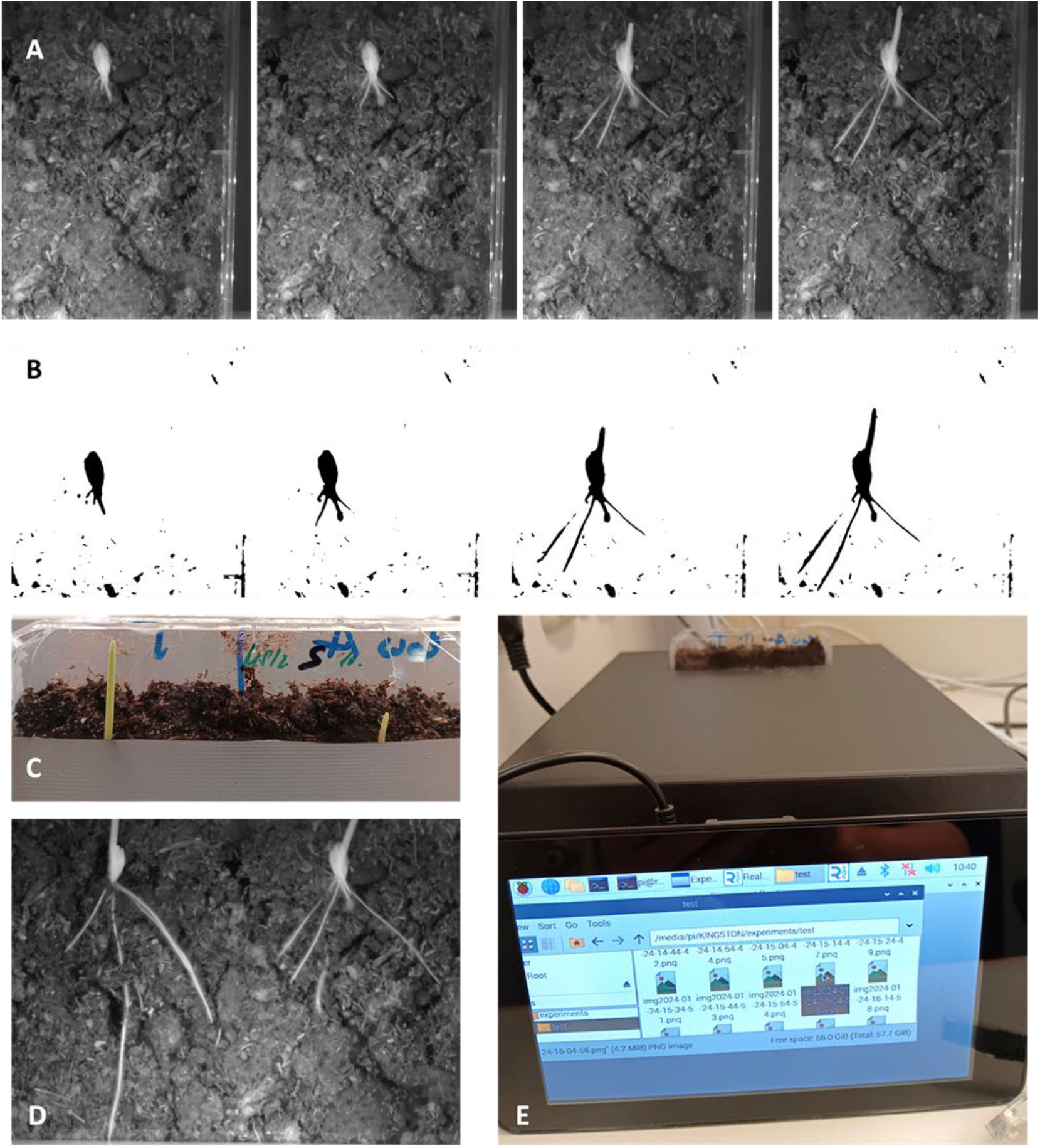
A) Selected images of barley seedlings are shown from a timeseries (acquired at 10-minute intervals and represent a 48-hour growth period). Simultaneous evaluation of root opening angle, seminal root number and dynamics of root growth rate is possible. Further traits that are easy to extract are mesocotyl growth rate and adaptation to different soil textures and compositions. B) The obtained images from the imaging chamber are suitable for batch segmentation with the RhizoVision Explorer (Seethepalli et al., 2021). The segmentation process is more effective when the soil is darker and humidity levels are low, which results in high contrasts levels between the barley root and the background. C) The image captures the emergence of barley coleoptiles. The imaging chamber allows the shoot to be exposed to light for long-term experiments. D) The image shows the roots at 5 days after germination grown in the imaging chamber. Images are acquired under few seconds IR light exposure (880nm). The roots correspond to the coleoptiles from panel C). E) The imaging setup allows continuous evaluation of the growth status of the germinating seedlings. The touch screen enables the selection of already taken images, which allows qualitative control of the experimental setup during the running experiment. Depending on the chosen dimensions of the imaging chamber, observation of growth dynamics over several days to weeks is possible to enable continuous root growth observation without disrupting seedling growth.

## 5. Summary

The aim of this paper is to introduce an updated imaging system that allows to monitor roots grown on soil and shaded from direct illumination, while letting the shoot develop in light. This system addresses the challenges of most root phenotyping methods that require exposing the roots to light, which can affect their morphology and physiology. With this system, we currently aim to dissect how different genotypes and environmental factors influence the number, angle, and elongation rate of barley seminal roots, which are crucial traits for cereal adaptation and productivity. Moreover, this imaging chamber can also be applied to monitor the dynamic establishment of root architecture of any other plant species, whether they are grown on soil, germination paper or agar plates and under different controllable growth conditions. Our imaging chamber is easy to construct, inexpensive, and suitable for high-throughput phenotyping experiments. It allows us to track root outgrowth from the onset of germination, if required also to investigate the growth dynamics of hypocotyl or mesocotyl, respectively.

The imaging chamber captures root images via a Raspberry Pi camera module at any desired interval to measure the development of selected root traits. We use infrared LEDs (880nm) as the light source for the camera, which do not trigger any detectable stress responses in the roots, according to previous studies. Our detailed step-to-step tutorial allows an easy and fast assembly of the imaging chamber, whereby the dimensions of the chamber and growth conditions can be adjusted to individual requirements regarding research object and scientific question. Furthermore, the advantages of the DDrC chamber include the possibility to track roots without any further modification of the plant by using IR LEDs. Compared to other methods, such as fluorescence or luminescence imaging, which require the use of chemical or genetic markers that may interfere with the root physiology or development. Moreover, our system can image the roots in a high-throughput manner, as it can process a large number of images in a short time. Also, due the inexpensive setup several DDrC chambers can be assembled and used in parallel. Although the method only allows to capture roots in a two-dimensional plane, as it uses a single camera that captures images from one angle, the obtained data inform about crucial root traits that are required for efficient adaptation of plant resilience. 3D methods such as X-ray CT or MRI, require longer scanning and reconstruction times, and more expensive equipment, and not everyone has access to the technology. We conclude that our system is a useful tool accessible for everyone to perform dynamic root phenotyping. Furthermore, we suggest to implement our system to explore the genetic and environmental factors that affect the root system architecture and development, and to identify the root traits that are associated with plant performance and adaptation. Finally, we also recommend that for future application the system can be improved by using updated versions of the camera module, other parts of the setup and especially image processing techniques, which happen to be adjusted continuously, to increase the resolution, depth, and accuracy of the root images.

## Supporting information

Supplement information describing data evaluation

