## Supplement information describing data evaluation for "Dynamic Dark Root Chamber – Advancing non-invasive phenotyping of roots kept in darkness using infrared imaging"

### Data acquisition and root trait analysis

Root exploration of the soil is essential for plant fitness and, in the case of cereals, for yield performance. Roots must reach water and nutrients as fast as possible, which requires dynamic adaptation of specific root traits. In drought conditions, cereal primary and seminal roots need to grow quickly and straight towards the potential groundwater, which demands a downward orientation and a narrow root opening angle (Maqbool *et al.*, 2022). These root traits have been linked to improved cereal drought resilience. Root structure and dynamics are partly determined by genetic factors, but they are also influenced by environmental factors that affect root shape, architecture, and function. We developed an imaging chamber to monitor root growth establishment continuously under controlled conditions from the moment of germination. This enables us to differentiate the role and growth behavior of the primary root, the radicle, in seedling establishment, from the growth dynamics of the first emerging seminal roots. Additionally, we can track the dynamic changes of root opening angle caused by mechanical interaction with the soil. The root growth dynamics and root angle can be measured in subsequent images manually with ImageJ/Fiji (Schindelin *et al.*, 2012), which yields high accuracy, but is time-consuming when several genotypes are investigated under different growth conditions (Fig. 1 A-C). Partially automated extraction of root traits is possible by applying RhizoVision Explorer (Seethepalli *et al.*, 2021) on the individual images and seedlings, which allows us to obtain information about root width and convex hull, which describe the occupancy and growth behavior across the soil levels (Fig. 2 A-C). Tracking total root length automatically is only feasible when the individual roots remain on the soil surface, otherwise, when they are partially covered by soil during the growth process, root trait evaluation is unreliable (Fig. 1 A).

Continuous acquisition of sequential images allows us to study fine responses of root growth over time. The root growth rate of the primary root, as well as the two first emerging seminal roots, is compared in Fig. 1 A between barley seedlings (genotype TISSA) grown in two independent experiments (Fig. 1 A and C). Root traits can be extracted from the moment of germination at most suitable timepoints for answering the individual scientific question and any chosen frequency, depending on the intervals of obtained images, without disruptive handling of the specimen (Fig. 1 B). Gaps in the measurement occur when the root tip cannot be temporarily located, for instance, when soil particles mask it, but over the whole course of the experiment the growth rate of the individual roots is clearly visible (Fig. 1 A and B). Moreover, root opening angle, which is formed by the outgrowing seminal roots, can be measured conveniently, and as the data show it fluctuates slightly over the chosen time course due to root-soil interactions (Fig. 1 B). The advantage of continuous imaging of the growth process of the emerging seminal roots stems from the unambiguous assignment of root outgrowth order. Timing of seminal root outgrowth, as well as their number and growth rate are crucial traits that underpin drought resilience, and can be analyzed in detail from images acquired in the Dynamic Dark Root imaging Chamber (DDrC).

**A, comparison of root growth dynamics between primary and seminal roots**

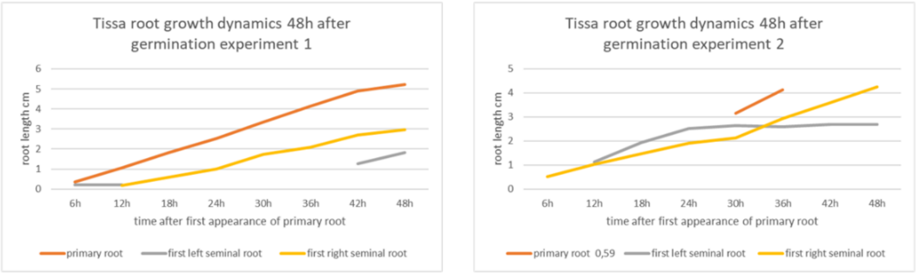


**B, raw data root length and opening angle**

| Tissa Experiment 1 | root length in cm | | | |
|---|---|---|---|---|
| | primary root | first left seminal root | first right seminal root | root opening angle |
| visible primary root | | | | |
| 6h | 0,351 | 0,197 | | |
| 12h | 1,056 | 0,215 | 0,181 | 78 |
| 18h | 1,814 | | 0,591 | 79 |
| 24h | 2,522 | | 1,009 | 78 |
| 30h | 3,337 | | 1,735 | 80 |
| 36h | 4,141 | | 2,092 | 77 |
| 42h | 4,91 | 1,268 | 2,697 | 80 |
| 48h | 5,221 | 1,824 | 2,954 | 83 |

| Tissa Experiment 2 | root length in cm | | | |
|---|---|---|---|---|
| | primary root | first left seminal root | first right seminal root | root opening angle |
| visible primary root | 0,59 | | | |
| 6h | | | 0,506 | |
| 12h | | 1,121 | 1,017 | 79 |
| 18h | | 1,923 | 1,462 | 81 |
| 24h | | 2,52 | 1,912 | 82 |
| 30h | 3,158 | 2,64 | 2,123 | 80 |
| 36h | 4,12 | 2,586 | 2,927 | 81 |
| 42h | | 2,687 | 3,588 | 80 |
| 48h | | 2,701 | 4,258 | 80 |

**C, evaluated roots**

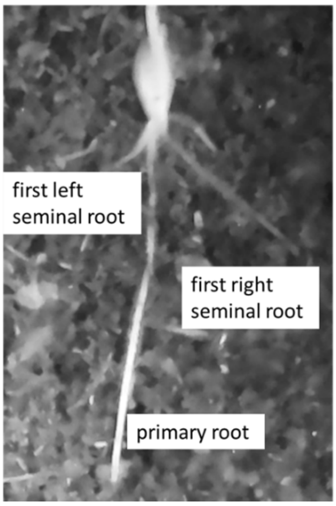


**Supplement Figure 1:** Evaluation of root length and seminal root opening angle using ImageJ.

To improve data acquisition efficiency, we applied automated segmentation and root trait extraction using RhizoVision Explorer (Fig. 2). The loose structure of the soil used in the experiments caused rearrangement of soil particles during root growth, which sometimes covered the root tip, making automated root length evaluation over the whole-time course difficult. Therefore, we focused on the evaluation of two other important root traits for plant fitness, maximum root width and convex hull. These two traits determine root system architecture distribution through the soil, which depends on the genetic background, but is also strongly influenced by environmental conditions, such as water availability, nutrient distribution and soil structure. To prevent non-root fragments from interfering with root segmentation, we chose a region of interest close to the root (Fig. 2 A). To enhance segmentation efficiency, we increased the contrast between root and soil by inverting the images directly in the RhizoVision Explorer application, and choosing threshold levels that filtered out most of the soil particles (Fig. 2 B). We analyzed maximum root width and convex hull from the moment of primary root visibility after germination and evaluated the growth behavior of the emerging seedling every 12h during 2 days after germination as an example (Fig. 2 C). This enabled us to evaluate the potential of root growth expansion and soil occupancy along with root growth direction at once, under controlled conditions, and to obtain information about dynamic root system architecture processes without disturbing plant growth, with the root protected from toxic direct illumination, but allowing the shoot to grow out of the DDrC.

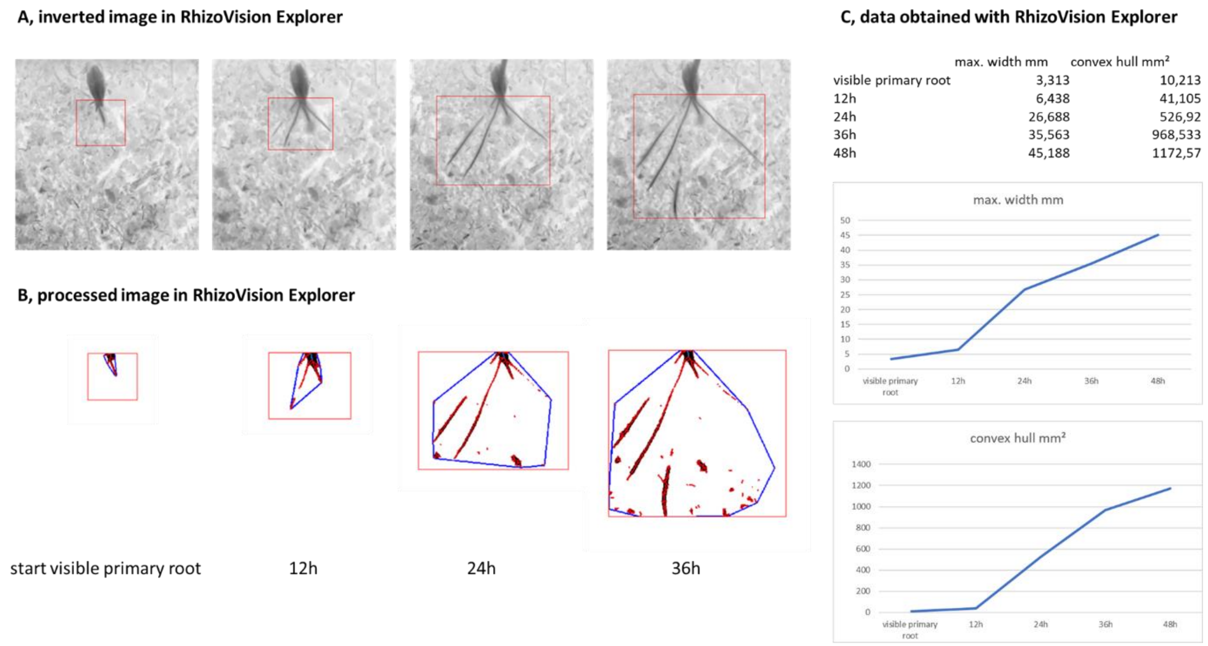


**Supplement Figure 2:** Evaluation of root maximal width and convex hull using RhizoVision Explorer.
